# Global Effects of Feature-based Attention Depend on Surprise

**DOI:** 10.1101/747204

**Authors:** Cooper A. Smout, Marta I. Garrido, Jason B. Mattingley

## Abstract

Recent studies have shown that prediction and attention can interact under various circumstances, suggesting that the two processes are based on interdependent neural mechanisms. In the visual modality, attention can be deployed to the location of a task-relevant stimulus (‘spatial attention’) or to a specific feature of the stimulus, such as colour or shape, irrespective of its location (‘feature-based attention’). Here we asked whether predictive processes are influenced by feature-based attention outside the current spatial focus of attention. Across two experiments, we recorded neural activity with electroencephalography (EEG) as human observers performed a feature-based attention task at fixation and ignored a stream of peripheral stimuli with predictable or surprising features. Central targets were defined by a single feature (colour or orientation) and differed in salience across the two experiments. Task-irrelevant peripheral patterns usually comprised one particular conjunction of features (standards), but occasionally deviated in one or both features (deviants). Consistent with previous studies, we found reliable effects of feature-based attention and prediction on neural responses to task-irrelevant patterns in both experiments. Crucially, we observed an interaction between prediction and feature-based attention in both experiments: the neural effect of feature-based attention was larger for surprising patterns than it was for predicted patterns. These findings suggest that global effects of feature-based attention depend on surprise, and are consistent with the idea that attention optimises the precision of predictions by modulating the gain of prediction errors.

**Significance Statement:** Two principal mechanisms facilitate the efficient processing of sensory information: *prediction* uses prior information to guide the interpretation of sensory events, whereas *attention* biases the processing of these events according to their behavioural relevance. A recent theory proposes to reconcile attention and prediction under a unifying framework, casting attention as a ‘precision optimisation’ mechanism that enhances the gain of prediction errors. Crucially, this theory suggests that attention and prediction interact to modulate neural responses, but this hypothesis remains to be tested with respect to feature-based attention mechanisms outside the spatial focus of attention. Here we show that global effects of feature-based attention are enhanced when stimuli possess surprising features, suggesting that feature-based attention and prediction are interdependent neural mechanisms.

## Introduction

Selective attention mechanisms enhance the processing of sensory stimuli that are relevant for guiding behaviour (Desimone & Duncan, 1995; Posner, 1994). Visual processing can be biased toward stimuli at a relevant location, commonly known as ‘spatial attention’, or toward stimuli that possess task-relevant features, known as ‘feature-based attention’ (Carrasco, 2011). Monkey neurophysiology studies (Martinez-Trujillo & Treue, 2004) and human neuroimaging studies (Gledhill, Grimsen, Fahle, & Wegener, 2015; Saenz, Buracas, & Boynton, 2002; Serences & Boynton, 2007) have demonstrated that neural responses to stimuli at task-irrelevant locations are enhanced when they possess task-relevant features, demonstrating that the effects of feature-based attention are global and dissociable from those of spatial attention. In humans, neural responses to visual stimuli at task-irrelevant locations can be enhanced when they possess surprising features (e.g., colour, orientation, motion), demonstrating that top-down ‘prediction’ mechanisms also exert a global effect on incoming sensory signals (Friston, 2005; Stefanics, Kremlácek, & Czigler, 2014). At present, it is unknown whether the global effects of visual feature-based attention can interact with those of prediction. Here we used electroencephalography (EEG) to measure neural responses to peripheral visual stimuli that were predictable or surprising along two feature dimensions (orientation and colour), and tested whether attending to a particular feature at fixation modulated the effect of prediction on neural responses to peripheral stimuli at task-irrelevant locations.

Predictive coding theories propose that top-down prediction signals effectively ‘silence’ bottom-up sensory signals that match the predicted content, leaving only the remaining *prediction error* to propagate forward and update a model of the sensory environment (Friston, 2005; Rao & Ballard, 1999). In addition to predicting the *content* of sensory signals, an optimal inference system should also estimate the level of uncertainty about its predictions (i.e., inverse precision; Hohwy, 2012). Recently, it has been proposed that selective attention mechanisms fulfil this role, optimising the expected precision of predictions by enhancing the activity of units encoding prediction errors for attended stimuli (Feldman & Friston, 2010; Friston, 2009, 2010). Recent studies have supported this theory by demonstrating that selective attention and prediction can interact under various circumstances (Auksztulewicz & Friston, 2015; Jiang, Summerfield, & Egner, 2013; Kok, Rahnev, Jehee, Lau, & De Lange, 2012; Marzecová, Widmann, SanMiguel, Kotz, & Schröger, 2017; Smout, Tang, Garrido, & Mattingley, 2019). However, selective attention mechanisms encompass distinct information-processing subcomponents (e.g., spatial attention, temporal attention) across sensory modalities (e.g., auditory, visual) and it is important to establish which of these subcomponents interacts with prediction and in what manner. In the visual domain, previous studies that reported an interaction between attention and prediction typically presented stimuli at task-relevant locations (Jiang et al., 2013; Kok, Rahnev, et al., 2012; Marzecová et al., 2017; Smout et al., 2019). One previous study found an effect of feature-based attention on mismatch responses to stimuli at task-irrelevant locations, but this study presented clearly visible targets that likely did not necessitate a tight focus of spatial attention on the central stimulus stream (Czigler & Sulykos, 2010). Thus, it remains unclear whether prediction can interact with global feature-based attention mechanisms that modulate neural responses to stimuli outside the spatial focus of attention.

Here we tested whether feature-based attention modulates the effect of prediction at task-irrelevant locations by comparing event-related potentials evoked by peripheral stimuli that either matched (‘congruent’) or mismatched (‘incongruent’) a cued feature of the target at fixation. Participants searched for targets at fixation while predictable or surprising task-irrelevant stimuli were presented in the periphery. We conducted two experiments that differed in the salience of central targets and distractors to investigate whether the strength of the top-down feature-set modulates the neural interaction between feature-based attention and prediction.

## Methods

### Experiment 1: Effects of feature-based attention at fixation on neural responses to predicted and surprising peripheral stimuli

#### Participants

Twenty-four healthy adults (13 female, 11 male, age = 22.08 ± 2.38 years) with normal or corrected-to-normal vision were recruited for participation via an online portal at The University of Queensland. The study was approved by The University of Queensland Human Research Ethics Committee, and all participants provided written, informed consent before commencing the experiment.

#### Stimuli and apparatus

Participants were positioned at a viewing distance of 57 cm and seated in a comfortable armchair in an electrically shielded laboratory. Stimuli were presented on a 61 cm LED monitor (Asus, VG248QE) with a 1920 × 1080 pixel resolution and refresh rate of 120 Hz, using PsychToolbox presentation software (Kleiner, M., Brainard, D., Pelli, D., Ingling, A., Murray, R., & Broussard, 2007) for Matlab (v.15b) running under Windows 7 with an NVidia Quadro K4000 graphics card. The intensity of the green phosphor was adjusted per participant to produce subjective equiluminance with that of the red phosphor at full intensity. The equiluminance point was determined prior to the experiment using minimum motion photometry (Anstis & Cavanagh, 1983), with intensity values determined by two interleaved adaptive staircases (1 up-1 down, stopping after 15 reversals).

Central and peripheral stimuli were sinusoidal Gabors (diameter: 4.72°, spatial frequency: 0.94 c/°, 100% contrast) with one of two orientations (tilted 45° clockwise or counterclockwise of vertical) and one of two colours (red or green). Central stimuli were superimposed over a red-green noise patch (diameter: 4.72) and onset every 700 - 1400 ms for 66.67 ms. Twenty percent (20%) of central stimuli were targets (approximately 28 targets and 114 distractors per block). Multi-element peripheral stimuli (‘patterns’) were arranged in three concentric circles (radii: 4.72°, 8.49°, 12.26°; containing 8, 14, and 20 Gabors, respectively; *Figure 1A*). Peripheral patterns were presented every 350 ms for 66.67 ms (428 events per block) on top of a background that alternated between uniform red and green pixels at the screen refresh rate (120 Hz), producing a subjective percept of a uniform brown background. During each block, peripheral patterns were more likely to contain one of the four possible feature conjunctions (e.g. clockwise-tilted red Gabors; 76% of presentations, *standards*), with the other three feature conjunctions being rare and of equal likelihood (8% each, *deviants*). Standards were pseudo-randomized across blocks, and the order of deviants was pseudo-randomized within blocks.

**Figure 1.**
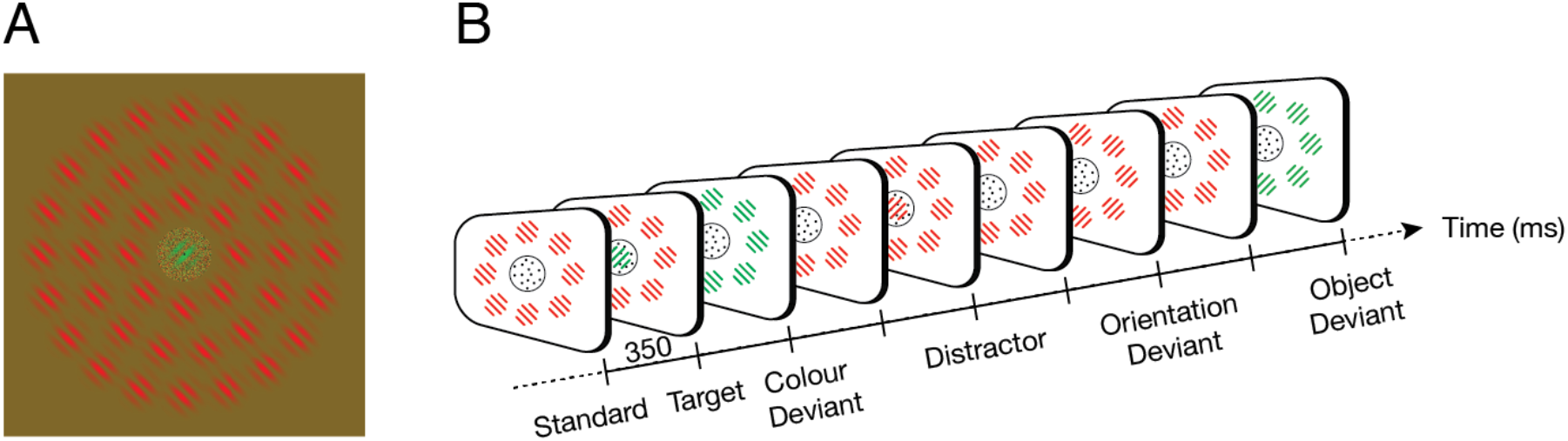
Stimulus display and task. (**A**) Stimulus display, showing a green clockwise-tilted target within the central noise patch and a peripheral pattern consisting of three concentric rings of red counterclockwise-tilted Gabors. (**B**) Simplified task diagram. In this example trial, participants monitored for green Gabors (targets) and ignored red Gabors (distractors) within the central noise patch. Peripheral patterns typically contained red counterclockwise-tilted Gabors (standards), but occasionally contained red clockwise-tilted Gabors (orientation deviant), green counterclockwise-tilted Gabors (colour deviant), or green clockwise-tilted Gabors (object deviant, i.e. deviating in both colour and orientation). In this example trial, the colour and object deviants shared features with the target (i.e., green) and would thus be labelled ‘congruent’.

#### Procedure

Participants were asked to fixate on a central dot and click a mouse button as soon as they detected a target in the stream of central stimuli, continuously throughout blocks (duration: 150 s), while ignoring central distractors and peripheral patterns (*Figure 1B*). In each block, Gabor targets were designated as either (1) red, (2) green, (3) clockwise-tilted, or (4) counterclockwise-tilted. Note that each condition dictated two of the four possible feature conjunctions as targets and two as distractors (e.g., if searching for clockwise-tilted targets, both red and green clockwise-tilted Gabors were valid targets).

Participants completed two practice blocks with auditory feedback after each response, before being fitted with the EEG cap and electrodes (see *EEG Data Acquisition*). Participants then completed 16 test blocks with target type and standard pattern features pseudorandomized across blocks (6848 peripheral patterns per session). Feedback on mean reaction time and the number of hits and false alarms was provided between blocks.

#### Behavioural Data Analysis

We investigated whether the feature-congruence and predictability of peripheral patterns affected participants’ detection of central targets. Targets were sorted into *prediction conditions* according to whether the preceding pattern (i.e., the peripheral stimulus presented up to 700 ms prior to peak target contrast) was a standard (‘predicted’) or a deviant (‘surprising’), and *feature-congruence conditions* according to whether the preceding peripheral pattern matched the features of the central target (‘congruent’) or distractor (‘incongruent’). Participant responses were scored as hits if they occurred within 1 s of the onset of a target. Successive responses within this window were ignored, as were any responses that occurred within 250 ms of a preceding response. Because we observed differences in hit rates and reaction times between target feature conditions (i.e., the feature that participants searched for at fixation, e.g. ‘red’), we first normalised hit rates and reaction times within each target feature condition, separately for feature-congruence and prediction conditions, and then collapsed across the target feature conditions. The resulting normalised hit rates and reaction times were then subjected to two-way repeated measures ANOVAs to assess the effects of peripheral pattern prediction (two levels: predicted, surprising) and feature-congruence (two levels: congruent, incongruent) on target detection.

#### EEG Data Acquisition

Participants were fitted with a 64 Ag-AgCl electrode EEG system (BioSemi Active Two: Amsterdam, Netherlands). Continuous data were recorded using BioSemi ActiView software (http://www.biosemi.com), and were digitized at a sample rate of 1024 Hz with 24-bit A/D conversion and a .01 – 208 Hz amplifier band pass. All scalp electrode offsets were adjusted to below 20μV prior to beginning the recording. Pairs of flat Ag-AgCl electro-oculographic electrodes were placed on the outside of both eyes, and above and below the left eye, to record horizontal and vertical eye movements, respectively.

#### EEG Preprocessing

EEG recordings were processed offline using the EEGlab toolbox in Matlab (Delorme & Makeig, 2004). Data were resampled to 256 Hz and high-pass filtered with a passband edge at 0.5 Hz (1691-point Hamming window, cut-off frequency: 0.25 Hz, −6 db). Raw data were inspected for the presence of faulty scalp electrodes (none were found). To clean the data, we applied an iterative process of artifactual epoch and component rejection using independent component analyses (ICA). The data were segmented into 350 ms epochs surrounding Gabor onsets (50 ms pre- and 300 ms post-stimulus) and baseline activity prior to stimulus onset was removed from each epoch. Epochs were subjected to ICA, and the SASICA plugin for EEGlab (Chaumon, Bishop, & Busch, 2015) was used to identify blink, saccade, and focal trial components. Epochs were rejected if they met any of the following criteria: (1) blink component activity greater than ±10 μV between −50 and 150ms; (2) saccade component activity greater ±5 μV between 0 and 350 ms; (3) focal component activity exceeding a joint probability threshold of ±7 SD (5.5% of epochs were removed due to blink, saccade, or focal activity). The remaining epochs were then subjected to ICA for a second time, and SASICA was used again to identify artifactual components.

For further analysis, the resampled raw data were band-pass filtered between 0.5 and 40 Hz (1691-point Hamming window, cut-off frequencies: 0.25 and 40.25 Hz, −6 db) and segmented into 550 ms epochs surrounding Gabor onsets (100 ms pre- and 450 ms post-stimulus). Epochs containing artefacts (identified previously using the first ICA) were removed. Independent component weights from the second ICA were applied to this new dataset and artefactual components (identified previously using the second ICA) were removed. Baseline activity in the 100 ms prior to stimulus onset was removed from each epoch.

#### Event-Related Potential and Bayes Factor Analyses

Peripheral patterns were sorted into prediction conditions based on whether they were standards (repeated at least 4 times; ‘predicted’) or deviants (‘surprising’), and attention conditions based on whether they shared features with central targets (‘congruent’) or distractors (‘incongruent’) in the central task. Trials in each attention and prediction condition were averaged within participants to produce event-related potentials (ERPs) for each individual. Statistical analyses of condition ERPs were conducted using two-tailed cluster-based permutation tests across participants (Monte-Carlo distribution with 5000 permutations, *p*_*cluster*_<0.05; sample statistic: dependent samples *t*-statistic, aggregated using the maximum sum of significant adjacent samples, *p*_*sample*_<.05) in the Fieldtrip toolbox for Matlab (Oostenveld, Fries, Maris, & Schoffelen, 2011). Statistical analyses of univariate condition averages were conducted using paired-samples *t*-tests and Bayesian analyses. The Bayes factor analyses allowed for quantification of evidence in favour of either the null or alternative hypothesis, with *BF*_*10*_ > 3 indicating substantial support for the alternative hypothesis and *BF*_*10*_ < 0.33 indicating substantial support for the null hypothesis. Bayes factors were computed using the Dienes (2014) calculator in Matlab.

### Experiment 2: Replication with individually thresholded manipulation of feature-based attention at fixation

In Experiment 1, the high contrast targets were detected at near-ceiling levels. To investigate whether the neural interaction between feature-based attention and prediction is sensitive to the strength of the top-down feature set, we conducted a second experiment in which central targets and distractors were individually thresholded to be less salient. Except for the minor methodological differences noted below, Experiment 2 was the same as Experiment 1 and thus afforded an opportunity to replicate the original results in a separate group of participants.

#### Methods

A new cohort of 24 healthy adults with normal (or corrected-to-normal) vision was recruited to participate in Experiment 2 (12 female, 12 male, age = 22.17 ± 2.88 years, mean ± SEM). The stimuli and apparatus were identical to those used in Experiment 1 (*Figure 1*), except that in Experiment 2 the central stimuli (targets and distractors) were presented at lower contrast and with a sinusoidal onset and offset profile (total duration: 700 ms). The peak contrast of the central stimuli was determined during the two practice blocks, using a transformed and weighted up/down adaptive staircase configured to approximate 83% detection of targets (up/down step ratio: 1/3, up/down size ratio: .1/.07; Garcı́a-Pérez, 1998). Blocks lasted for 150 s (as per Experiment 1) for all except two participants, for whom blocks lasted for 120 s (due to time constraints for these two individuals). Participant responses were scored as hits if they occurred within 1 s of the peak target contrast (i.e., within 1.35 s of target onset, accounting for the 350 ms on-ramp). During EEG preprocessing, we interpolated 11 faulty electrodes (across 5 participants) using the average activation across neighbouring electrodes (defined by the EEGlab Biosemi 64 template) and removed 4.1% of epochs due to blink, saccade, or focal component contamination.

## Results

### Experiment 1: Effects of feature-based attention at fixation on neural responses to predicted and surprising peripheral stimuli

#### Feature-Congruent Peripheral Patterns Interfere with Target Detection at Fixation

We first asked whether the congruence between peripheral pattern and central target features affected participants’ detection of central targets shortly after pattern onset. There was no significant main effect of feature-congruence on normalized hit rates (congruent = 94.41 ± 1.28%, 0.06 ± 0.19 z-normalised, incongruent = 93.63 ± 1.39%, −0.08 ± 0.20 z-normalised, *F*(1,23) = 3.33, *p* = .081, *η*_*p*_^*2*^ = .013). There was a significant main effect of feature-congruence on normalised reaction times, however, with participants responding more slowly to central targets preceded by congruent peripheral patterns (438.80 ± 9.43 ms, mean ± SEM; 0.04 ± 0.19 z-normalised) than to those preceded by incongruent peripheral patterns (436.09 ± 9.53 ms; −0.04 ± 0.20 z-normalised, *F*(1,23) = 5.70, *p* = .026, *η*_*p*_^*2*^ = 0.20). This finding suggests that participants were more distracted by peripheral patterns with task-relevant (congruent) features, relative to those with task-irrelevant (incongruent) features, and is consistent with the theory that involuntary orienting to task-irrelevant stimuli is contingent on attentional control settings (Folk, Remington, & Johnston, 1992).

#### Peripheral Pattern Prediction Does Not Affect Target Detection

In a second analysis we asked whether the predictability of peripheral patterns affected behavioural responses to subsequent central targets. There was no significant effect of peripheral pattern prediction on normalised hit rates (predicted = 94.39 ± 1.22%, 0.04 ± 0.18, surprising = 93.65 ± 1.47%, −0.06 ± 0.21, *F*(1,23) = 1.63, *p* = .215, *η*_*p*_^*2*^ = .07) or normalized reaction times (predicted = 437.12 ± 9.66 ms, −0.01 ± 0.20, surprising = 437.76 ± 9.32 ms, 0.01 ± 0.19, *F*(1,23) = 0.27, *p* = .605, *η*_*p*_^*2*^ = .01), and no interaction between prediction and feature-congruence on either normalised hit rates (*F*(1,23) = 0.31, *p* = .582, *η*_*p*_^*2*^ = .01) or normalised reaction times (*F*(1,23) = 0.15, *p* = .701, *η*_*p*_^*2*^ = .01). These findings suggest that the predictability of peripheral patterns did not modulate the extent to which participants were distracted from their task at fixation.

#### Prediction Decreases Neural Activity

We next assessed the main effect of prediction on neural activity by comparing ERPs to peripheral deviant patterns (‘surprising’ patterns, collapsed across orientation, colour, and object deviants) and standard patterns that had been repeated at least 4 times (‘predicted’ patterns). Relative to baseline, standards evoked smaller neural responses than deviants (*Figure 2*). Over posterior electrodes, the neural response to standards was significantly reduced relative to deviants during both the early negative deflection (i.e. standards > deviants; 82 - 164 ms, *p* = .020) and the late positive deflection (i.e. deviants > standards < deviants; 242 – 348 ms, *p* = .010; *Figure 2B*). Over frontal electrodes, the neural response to standards was significantly reduced relative to deviants during both the early positive deflection (i.e. standards < deviants; 90 - 230 ms, *p* < .001) and the late negative deflection (i.e. standards > deviants; 254 – 348 ms, *p* = .008; *Figure 2A*). These effects are consistent with the theory that surprising stimuli (deviants) produce greater prediction errors than predicted stimuli (standards; Friston, 2005, 2009; Rao & Ballard, 1999).

**Figure 2.**
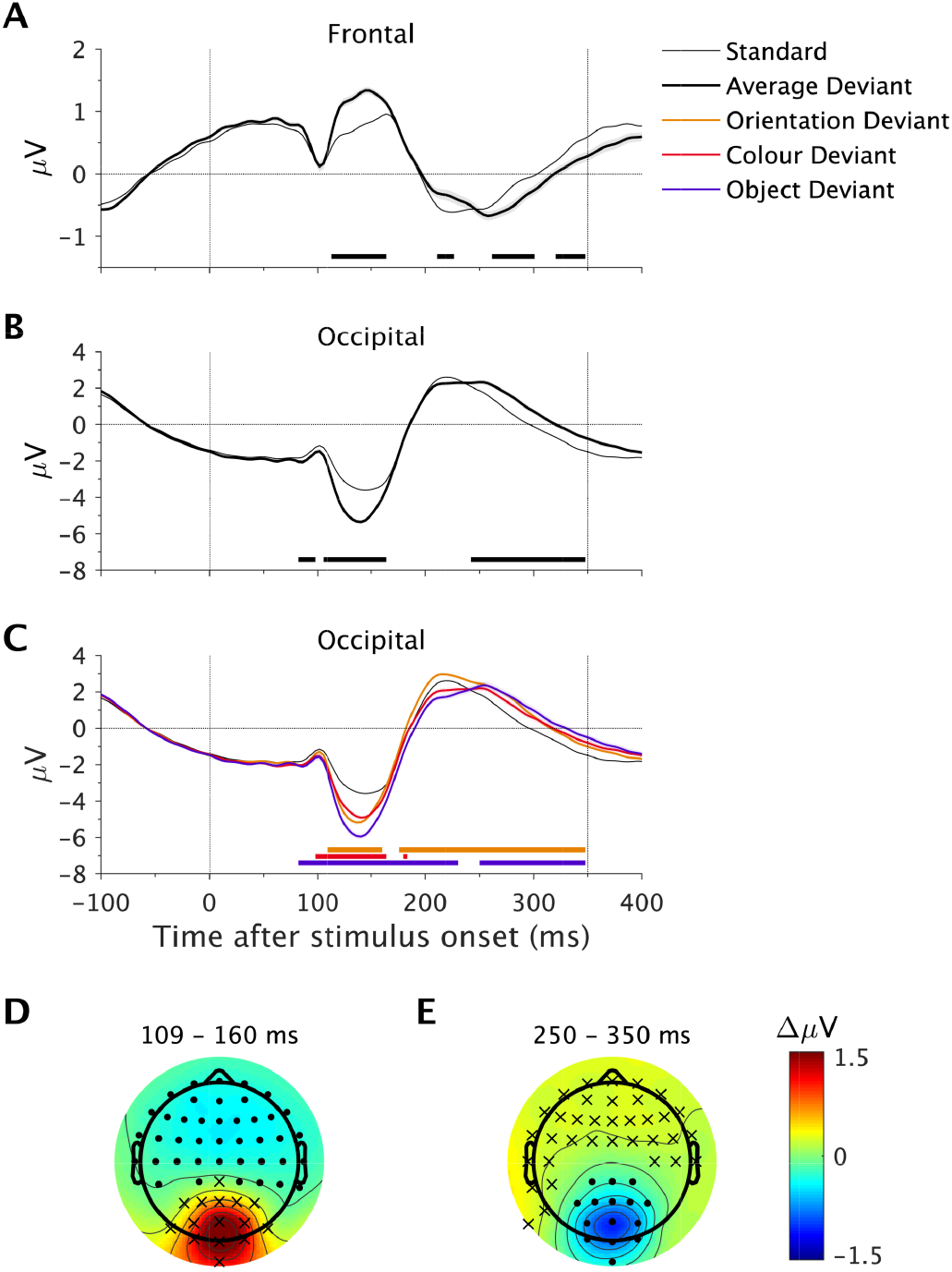
Main effect of prediction in Experiment 1. (**A-B**) ERPs evoked by standards and deviants (collapsed across deviant types) at frontal electrodes (Fz, F1, F3, AFz, AF3, AF4; **A**) and occipital electrodes (Oz, O1, O2, POz, PO3, PO4; **B**). Shading indicates the within-subject standard error of the mean, calculated relative to standards. Black bars along the x-axis denote significant timepoints at the displayed electrodes (cluster-corrected). (**C**) ERPs evoked by standards and each of the three deviant conditions. Shading indicates the within-subject standard error of the mean, calculated separately for each deviant condition relative to standards. Yellow, red and purple bars along the x-axis denote significant differences between standards and each corresponding deviant condition (cluster-corrected). (**D-E**) Headmaps show the effect of prediction (standard minus average deviant) during the indicated time windows. Asterisks and dots denote electrodes with larger or smaller responses, respectively, across at least 33% of the averaged time points (cluster-corrected).

We followed up this result with direct comparisons between standards and each type of deviant, which revealed similar effects to those reported above for the average deviant condition (*Figure 2C*). Early posterior negativities were smaller in response to standards than orientation deviants (109 – 160 ms, *p* = .033), colour deviants (98 – 164 ms, *p* = .040), and object deviants (82 – 348 ms, *p* < .001), and late posterior positivities were significantly smaller in response to standards than orientation deviants (176 – 348 ms, *p* < .001) and object deviants (250 – 348 ms, *p* = .019). Early frontal positivities were smaller in response to standards than orientation deviants (98 – 164 ms, *p* = .002), colour deviants (86 – 238 ms, *p* = .001), and object deviants (102 – 238 ms, *p* < .001), and late frontal negativities were smaller in response to standards than orientation deviants (242 – 348 ms, *p* < .001) and object deviants (84 – 348 ms, *p* < .001).

#### Visual Mismatch Negativities Are Additive Across Feature Deviations

Because previous investigations have suggested that the visual mismatch negativity (vMMN) is non-additive across feature deviations (Czigler & Sulykos, 2010), we also tested for differences between vMMNs evoked by each type of deviant (orientation, colour, or object). We used a data-driven approach to identify spatiotemporal samples (electrodes × timepoints) that were significantly different from standards in all three deviant conditions (electrodes: Pz, P1, P2, P3, P4, POz, PO3, PO4, Oz, O1, O2, Iz; timepoints: 109 – 160 ms) and then averaged across these samples to produce one amplitude value per deviant condition and participant. We then compared each pair of deviant conditions with paired-samples *t*-tests and Bayesian analyses, using a uniform prior with upper and lower bounds set to the average vMMN amplitude. As can be seen in *Figure 2C*, there was no difference between the orientation (−1.02 ± 0.13 μV) and colour vMMN (−0.98 ±Z0.14 μV, *t*(23) = −0.35, *p* = .733, *BF_10_* = 0.14). In contrast, the object vMMN (−1.64 ± 0.16 μV) was significantly larger than both the orientation vMMN (*t*(23) = −5.39, *p* < .001, *BF_10_* = 2.4 × 10^5^) and the colour vMMN (*t*(23) = −7.06, *p* < .001, *BF_10_* = 6.3 × 10^9^), suggesting that the vMMN is sensitive to features of the deviant stimulus.

#### Feature-based Attention Decreases Neural Activity

We assessed the main effect of feature-based attention by comparing ERPs to peripheral patterns that shared features with targets (‘congruent’) or distractors (‘incongruent’) in the central detection task. Congruent peripheral patterns evoked a smaller positivity over posterior electrodes than incongruent patterns late in the epoch (188 – 305 ms, *p* = .004; *Figure 3B,D*). This effect was matched by a polarity-reversed activity profile over frontal electrodes (191 – 309 ms, *p* = .003; *Figure 3A,D*).

**Figure 3.**
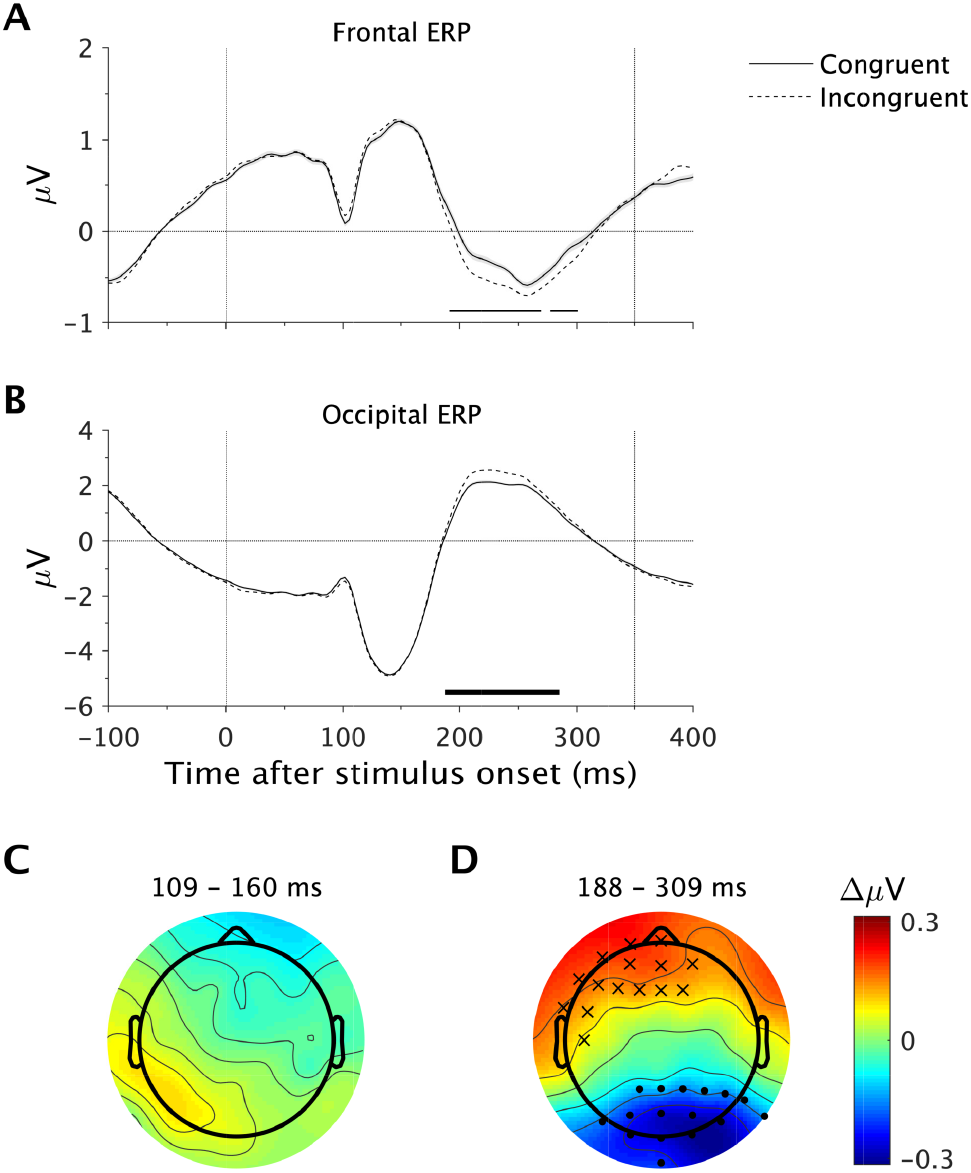
Main effect of feature-based attention in Experiment 1. (**A-B**) Congruent and incongruent ERPs are collapsed across prediction conditions separately for frontal electrodes (Fz, F1, F3, AFz, AF3, AF4; **A**) and occipital electrodes (Oz, O1, O2, POz, PO3, PO4; **B**). Shading indicates the within-subject standard error of the mean. Black bars along the x-axis denote significant differences at the displayed electrodes (cluster-corrected). (**C-D**) Headmaps show the effects of feature-based attention (congruent minus incongruent) during the indicated time windows. Asterisks and dots denote electrodes with larger, or smaller responses, respectively, in at least 33% of the averaged time points (cluster-corrected).

#### The Effect of Feature-based Attention Depends on Surprise

Next, we investigated the interaction between feature-based attention and prediction by subtracting the standard ERP from the deviant ERP (i.e., the mismatch response, collapsed across deviant conditions) and comparing difference waves between congruent and incongruent conditions (*Figure 4*). Over posterior electrodes, the mismatch response was more negative for congruent stimuli than for incongruent stimuli late in the epoch (203 – 285 ms, *p* = .041; *Figure 4B*). Inspection of individual condition ERPs (*Figure 4E*) revealed that the significant interaction was driven by a larger (negative) effect of feature-based attention on the neural response to deviants, relative to standards.

**Figure 4.**
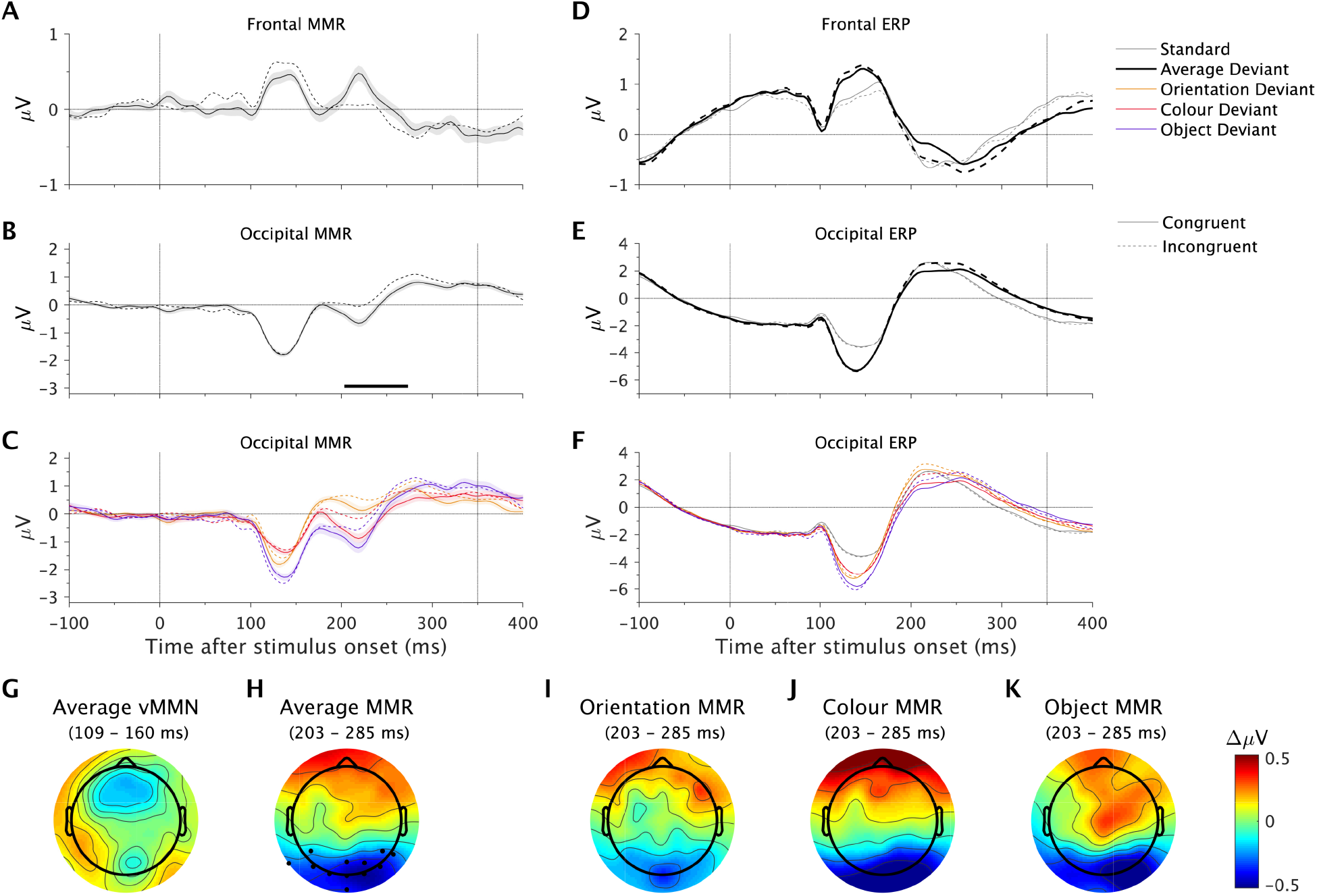
Interaction between feature-based attention and prediction in Experiment 1. (**A-B**) Average mismatch response (MMR; average deviant minus standard) collapsed across frontal electrodes (Fz, F1, F3, AFz, AF3, AF4; **A**) and occipital electrodes (Oz, O1, O2, POz, PO3, PO4; **B**). Solid lines represent the congruent condition and dotted lines represent the incongruent condition. Shading indicates the within-subject standard error of the mean. The black bar along the x-axis denotes significant differences at the displayed electrodes (cluster-corrected). (**C**) Mismatch responses at occipital electrodes for individual deviant conditions. (**D-E**) ERPs evoked by standards and deviants (averaged across deviant types), shown separately for congruent (solid) and incongruent (dotted) conditions. (**F**) ERPs for individual deviant conditions, shown separately for congruent (solid) and incongruent (dotted) conditions. (**G-H**) Headmaps show the effect of feature-based attention (congruent minus incongruent) on the average deviant mismatch response (average deviant minus standard) during the early vMMN (**G**) and late interaction time windows (**H**). Dots denote electrodes with significant differences in at least 33% of the averaged time points (cluster-corrected). (**I-K**) Effect of feature-based attention (congruent minus incongruent) on the orientation mismatch response (**I**), colour mismatch response (**J**) and object mismatch response (**K**) during the late interaction time window. Note that cluster-based permutation tests were not conducted on these differences.

We followed up this finding by averaging spatiotemporal samples spanned by the significant effect, separately for each deviant condition. We then compared congruent and incongruent conditions with paired *t*-tests and Bayesian analyses (uniform prior with upper and lower bounds set to the average amplitude across all conditions). Feature-based attention decreased the mismatch response to all three deviant types (orientation: congruent = 0.13 ± 0.10 μV, incongruent = 0.42 ± 0.13 μV, *t*(23) = −2.16, *p* = .041, *BF*_10_ = 1.18; colour: congruent = −0.45 ± 0.17 μV, incongruent = 0.05 ± 0.14 μV, *t*(23) = −3.28, *p* = .003, *BF*_10_ = 1.41; object: congruent = −0.53 ± 0.19 μV, incongruent = −0.06 ± 0.14 μV, *t*(23) = −2.92, *p* = .008, *BF*_10_ = 1.28; *Figure 4C*). These findings suggest that feature-based attention modulates the effect of prediction on neural responses to stimuli at task-irrelevant locations, irrespective of the predicted feature (or combination of features).

#### The Visual Mismatch Negativity is Not Modulated by Feature-based Attention

Because previous literature has provided evidence for an effect of feature-based attention on the vMMN (Czigler & Sulykos, 2010), we also used Bayes analyses to test for differences between congruent and incongruent conditions during the (non-significant) vMMN time period. Spatiotemporal samples spanning the common vMMN window (electrodes: Pz, P1, P2, P3, P4, POz, PO3, PO4, Oz, O1, O2, Iz; timepoints: 109 – 160 ms) were averaged to produce one amplitude value for each condition within participants. Congruent and incongruent conditions were compared within deviant conditions using paired-samples *t*-tests and Bayes analyses (uniform prior with upper and lower bounds set to the average amplitude across conditions). We found no difference between congruent and incongruent vMMNs for any deviant type (orientation: congruent = −1.09 ± 0.13 μV, incongruent = −0.95 ± 0.16 μV, *t*(23) = −1.23, *p* = .231, *BF*_10_ = .26; colour: congruent = −1.00 ± 0.16 μV, incongruent = −0.96 ± 0.14 μV, *t*(23) = −0.37, *p* = .713, *BF*_10_ = 0.13; object: congruent = −1.55 ± 0.19 μV, incongruent = −1.73 ± 0.15 μV, *t*(23) = 1.29, *p* = .208, *BF*_10_ = 0.33; *Figure 4C*).

Taken together, the results from Experiment 1 suggest that feature-based attention modulates the neural effect of prediction on neural responses to stimuli at task-irrelevant locations. This interaction emerged after (but not during) the vMMN time period for all deviant types, from approximately 200 ms after stimulus onset. We also found that the detection of high contrast targets at fixation was slower following peripheral patterns with target features, relative to those with distractor features, suggesting that feature-congruent peripheral patterns ‘captured’ attention to their location (Folk et al., 1992). Because our principle question of interest pertained to the neural interaction between feature-based attention and prediction *outside* the current spatial focus of attention, we conducted a second study in which target contrast was individually titrated for each participant to increase the task difficulty and ensure that attention remained fixed on the central target stream.

### Experiment 2: Replication with individually thresholded manipulation of feature-based attention at fixation

#### Peripheral Patterns Do Not Modulate Behaviour in a Demanding Feature-based Attention Task

In contrast to Experiment 1, there was no significant effect of feature-congruence on normalised reaction times in Experiment 2 (congruent: 391.79 ± 11.12 ms, −0.01 ± 0.18 z-normalised; incongruent: 392.49 ± 11.56 ms, 0.02 ± 0.18 z-normalised; *F*(1,23) = 1.00, *p* = .329, *η*_*p*_^*2*^ = .04), suggesting that the top-down feature set modulates the effect of congruent patterns on target detection and that the more difficult task in Experiment 2 contained spatial attention to the central target stream. In line with Experiment 1, all other behavioural effects were non-significant. Thus, there was no significant effect of feature-congruence on normalised hit rates (congruent = 74.90 ± 2.05%, −0.02 ± 0.16 z-normalised, incongruent = 75.23 ± 2.14%, 0.01 ± 0.17 z-normalised, *F*(1,23) = 0.37, *p* = .547, *η*_*p*_^*2*^ = .02). In addition, there was no significant effect of pattern prediction on normalised hit rates (predicted = 74.25 ± 2.07%, −0.06 ± 0.16 z-normalised, surprising = 75.88 ± 2.13%, 0.06 ± 0.17 z-normalised, *F*(1,23) = 3.64, *p* = .069, *η*_*p*_^*2*^ = 14) or on normalised reaction times (predicted = 388.79 ± 11.11 ms, −0.03 ± 0.18 z-normalised, surprising = 395.49 ± 11.58 ms, 0.04 ± 0.18 z-normalised, *F*(1,23) = 2.46, *p* = .130, *η*_*p*_^*2*^ = .10). Finally, there was no interaction between prediction and feature-congruence on either normalised hit rates (*F*(1,23) = 2.11, *p* = .160, *η*_*p*_^*2*^ = .08) or normalised reaction times (*F*(1,23) = 1.50, *p* = .233, *η*_*p*_^*2*^ = .06).

#### The Neural Interaction Between Feature-based Attention and Prediction Replicates With a Demanding Feature-based Attention Set

The neural effects observed in Experiment 2 (see *Figures 5-7*) were highly similar to those in Experiment 1 (see *Figures 2-4*). Prediction again modulated neural responses over posterior electrodes early (standards > deviants; from 78 ms, *p* < .001) and late (standards < deviants; 246 – 348 ms, *p* = .014) in the epoch (*Figure 5B*), with opposite early (standards < deviants; 74 - 238 ms, *p* < .001) and late effects (standards > deviants; prior to 348 ms, *p* < .001) over frontal electrodes (*Figure 5A*). Follow-up comparisons revealed similar effects of prediction on each deviant type (*Figure 5C*). Over posterior electrodes, standards evoked smaller early negativities than all deviant types (orientation deviants: 109 – 160 ms, *p* = .037; colour deviants: 90 – 348 ms, *p* < .001; object deviants; 78 – 348 ms, *p* < .001) and smaller late positivities than all deviant types (orientation deviants: 227 – 348 ms, *p* = .001; colour deviants; 250 – 348 ms, *p* = .029; object deviants: 250 – 348 ms, *p* = .022). Over frontal electrodes, standards evoked smaller early positivities than all deviant types (orientation: 47 – 156 ms, *p* = .0012; colour: 98 – 242 ms, *p* = .002; object: 78 – 242 ms, *p* < .001) and smaller late negativities than orientation deviants (234 – 348 ms, *p* < .001). As in Experiment 1, the vMMN was sensitive to features of the deviant stimulus (*Figure 5C*), with object deviants evoking a significantly larger vMMN (−1.86 ± 0.27 μV) than orientation deviants (−1.10 ± 0.16 μV, *t*(23) = −5.40, *p* < .001, *BF_10_* = 288,942.02) and colour deviants (−1.05 ± 0.19 μV, *t*(23) = −6.17, *p* < .001, *BF_10_* = 22,207,026.78). Again, there was no difference between orientation and colour vMMNs (*t*(23) = −0.42, *p* = .679, *BF_10_* = 0.12).

**Figure 5.**
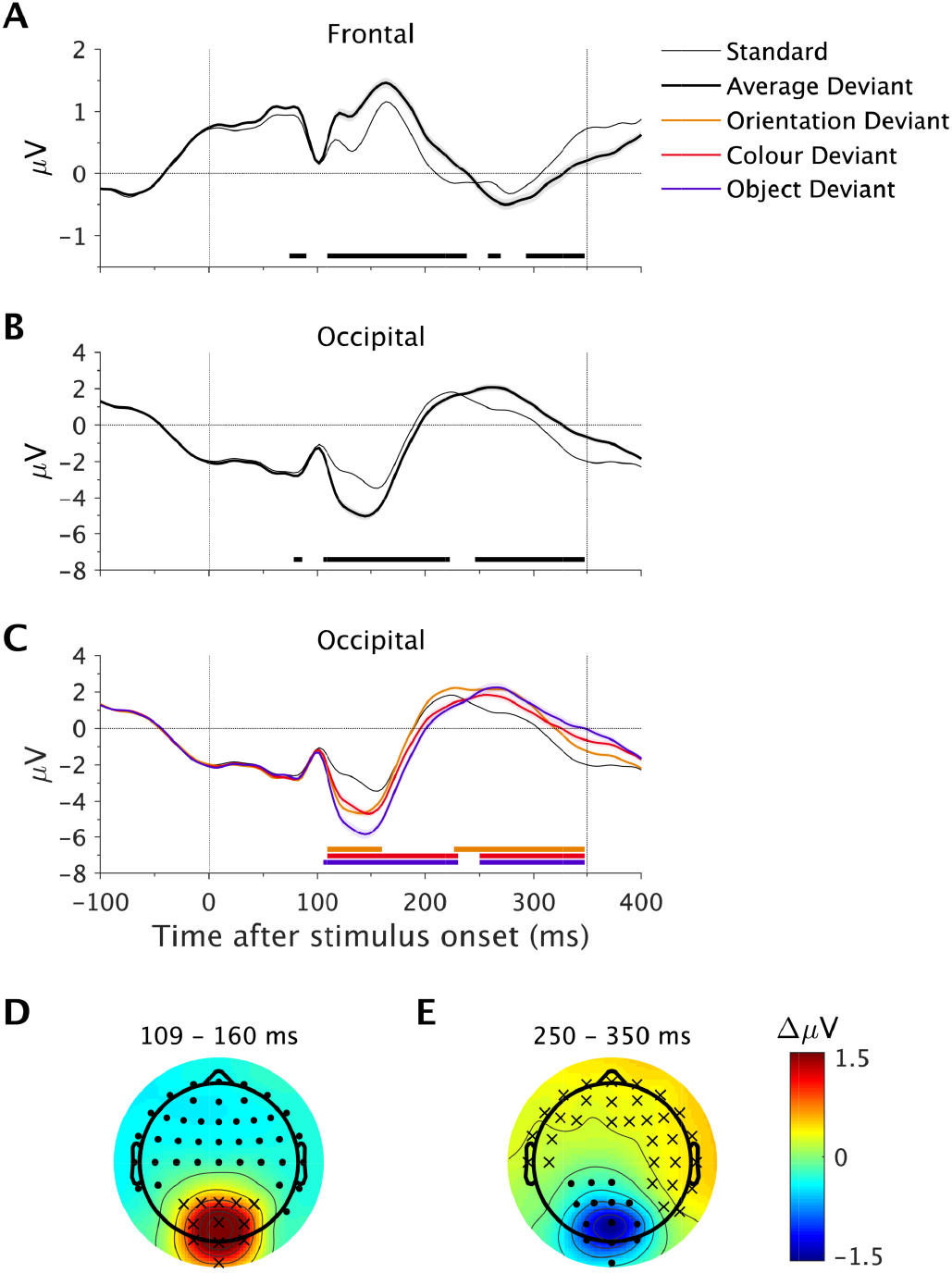
Main effect of prediction in Experiment 2. (**A-B**) ERPs evoked by standards and deviants (collapsed across deviant types) at frontal electrodes (Fz, F1, F3, AFz, AF3, AF4; **A**) and occipital electrodes (Oz, O1, O2, POz, PO3, PO4; **B**). Shading indicates the within-subject standard error of the mean, calculated relative to standards. Black bars along the x-axis denote significant timepoints at the displayed electrodes (cluster-corrected). (**C**) ERPs evoked by standards and each of the three deviant conditions. Shading indicates the within-subject standard error of the mean, calculated separately for each deviant condition relative to standards. Yellow, red and purple bars along the x-axis denote significant differences between standards and each corresponding deviant condition (cluster-corrected). (**D-E**) Headmaps show the effect of prediction (standard minus average deviant) during the indicated time windows. Asterisks and dots denote electrodes with larger, or smaller responses, respectively, across at least 33% of the averaged time points (cluster-corrected).

As in Experiment 1, congruent peripheral patterns evoked smaller positivities over posterior electrodes than incongruent patterns late in the epoch (195 – 273 ms, *p* = .045; *Figure 6B*). However, the polarity-reversed frontal effect observed in Experiment 1 was not significant in Experiment 2 (230 – 281 ms, *p* = .127).

**Figure 6.**
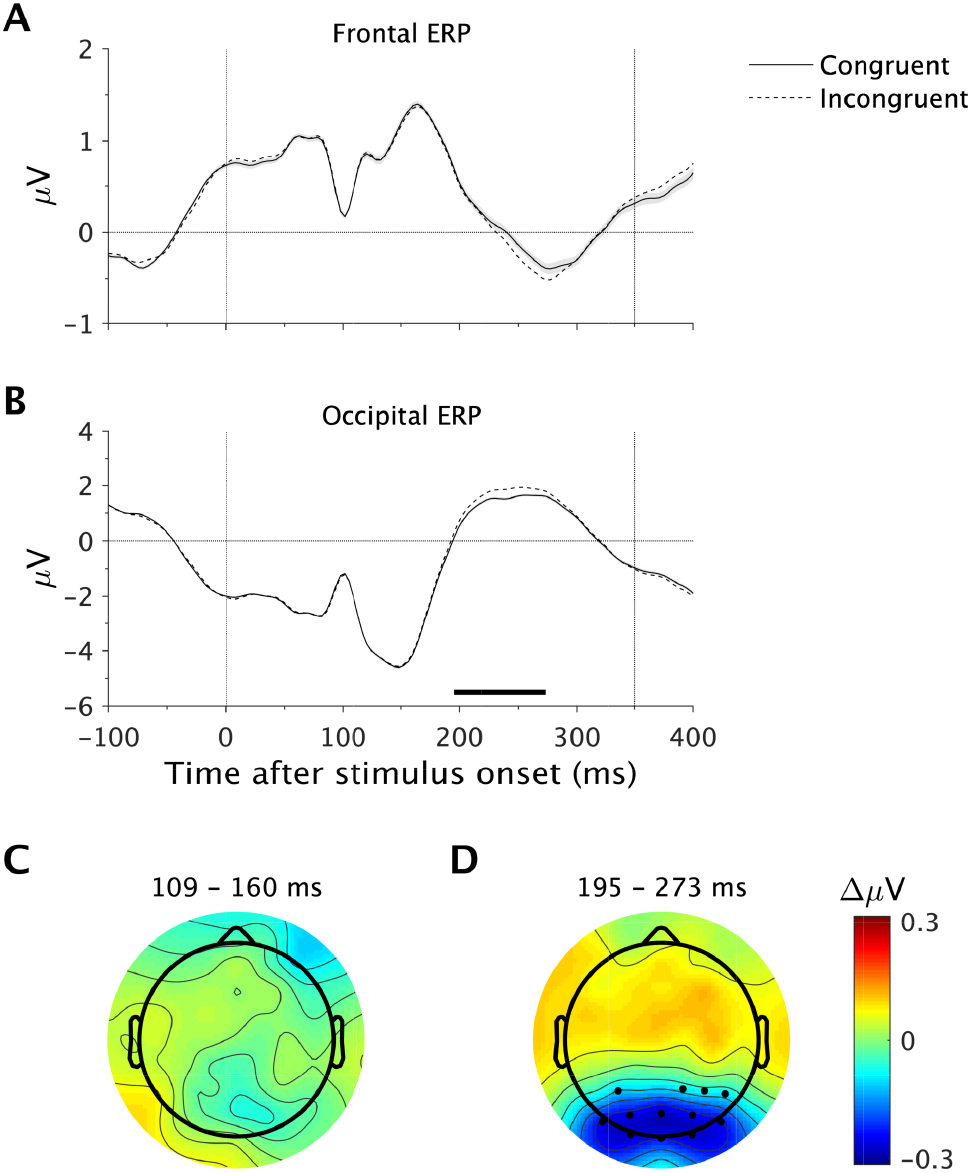
Main effect of feature-based attention in Experiment 2. (**A-B**) Congruent and incongruent ERPs are collapsed across prediction conditions separately for frontal electrodes (Fz, F1, F3, AFz, AF3, AF4; **A**) and occipital electrodes (Oz, O1, O2, POz, PO3, PO4; **B**). Shading indicates the within-subject standard error of the mean. The black bar along the x-axis denotes significant differences at the displayed electrodes (cluster-corrected). (**C-D**) Headmaps show the effects of feature-based attention (congruent minus incongruent) during the indicated time windows. Dots denote electrodes with smaller responses in at least 33% of the averaged time points (cluster-corrected).

Crucially, we replicated the significant interaction between feature-based attention and prediction observed in Experiment 1 (*Figure 7*). Congruent mismatch responses (deviants minus standards) were significantly smaller than incongruent mismatch responses over posterior electrodes late in the epoch (242 – 320 ms, *p* = .048; *Figure 7B,H*). We also observed an additional polarity-reversed effect over frontal electrodes that was absent in Experiment 1 (254 – 324 ms, *p* = .026; *Figure 7A,H*). Follow-up analyses revealed that feature-based attention significantly decreased the mismatch response to colour deviants (congruent = −0.01 ± 0.17 μV, incongruent = 0.31 ± 0.13 μV, *t*(23) = −3.52, *p* = .002, *BF*_10_ = 94.83, *Figure 7J*) and object deviants (congruent = 0.10 ± 0.21 μV, incongruent = 0.61 ± 0.14 μV, *t*(23) = −3.66, *p* =.001, *BF*_10_ = 51.42, *Figure 7K*) but only trended in the same direction for orientation deviants (congruent = 0.40 ± 0.11 μV, incongruent = 0.60 ± 0.10 μV, *t*(23) = −1.96, *p* = .062, *BF*_10_ = 2.35, *Figure 7I*). Again, we found no effect of feature-based attention on the vMMN evoked by any type of deviant (orientation: congruent = −1.09 ± 0.19 μV, incongruent = −1.11 ± 0.15 μV, *t*(23) = .18, *p* = .857, *BF*_10_ = .12; colour: congruent = −1.08 ± 0.19 μV, incongruent = −1.02 ± 0.22 μV, *t*(23) = −0.42, *p* = .676, *BF*_10_ = .15; object: congruent = −1.96 ± 0.28 μV, incongruent = −1.76 ± 0.27 μV, *t*(23) = −1.67, *p* = .11, *BF*_10_ = 0.43; *Figure 7C*).

**Figure 7.**
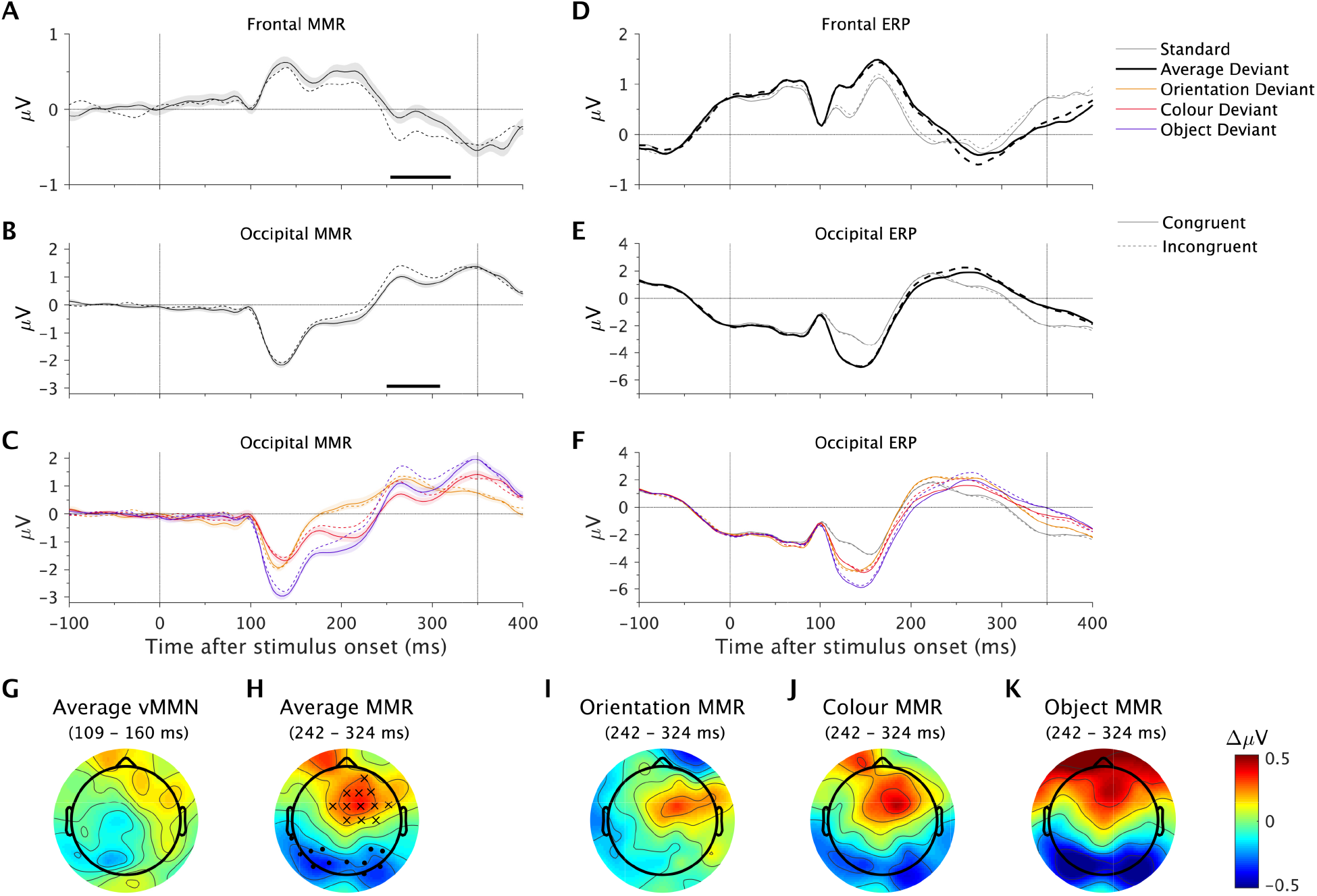
Interaction between feature-based attention and prediction in Experiment 2. (**A-B**) Average mismatch response (MMR; average deviant minus standard) collapsed across frontal electrodes (Fz, F1, F3, AFz, AF3, AF4; **A**) and occipital electrodes (Oz, O1, O2, POz, PO3, PO4; **B**). Solid lines represent the congruent condition and dotted lines represent the incongruent condition. Shading indicates the within-subject standard error of the mean. The black bar along the x-axis denotes significant differences at the displayed electrodes (cluster-corrected). (**C**) Mismatch responses at occipital electrodes for individual deviant conditions. (**D-E**) ERPs evoked by standards and deviants (averaged across deviant types), shown separately for congruent (solid) and incongruent (dotted) conditions. (**F**) ERPs for individual deviant conditions, shown separately for congruent (solid) and incongruent (dotted) conditions. (**G-H**) Headmaps show the effect of feature-based attention (congruent minus incongruent) on the average deviant mismatch response (average deviant minus standard) during the early vMMN (**G**) and late interaction time windows (**H**). Asterisks and dots denote electrodes with larger, or smaller responses, respectively, in at least 33% of the averaged time points (cluster-corrected). (**I-K**) Effect of feature-based attention (congruent minus incongruent) on the orientation mismatch response (**I**), colour mismatch response (**J**) and object mismatch response (**K**) during the late interaction time window. Note that cluster-based permutation tests were not conducted on these differences.

Overall, the findings from Experiment 2 replicate those from Experiment 1 to show that feature-based attention and prediction interact in their modulation of neural responses to stimuli at task-irrelevant locations. Similar to Experiment 1, this interaction emerged after (but not during) the vMMN time period for all deviant types, from approximately 240 ms after stimulus onset.

## Discussion

Here we investigated whether prediction interacts with feature-based attention outside the spatial focus of attention. To achieve this, we measured neural responses to surprising and predicted stimuli – deviants and standards, respectively – presented at task-irrelevant locations. Task-irrelevant peripheral patterns shared features with either the targets (congruent) or distractors (incongruent) in a central search task. Across two experiments, we replicated the finding that feature-based attention decreased neural responses to surprising but not predicted task-irrelevant stimuli in the periphery of vision. This finding suggests that the global neural mechanisms of feature-based attention and prediction are interdependent, and supports the theory that attention increases the gain of prediction errors (Feldman & Friston, 2010).

Consistent with previous literature, prediction modulated early and late neural responses to stimuli in both experiments (*Figures 2 & 5*). Early responses (approximately 100 to 160 ms) over posterior electrodes were more negative for surprising stimuli than predicted stimuli, consistent with the commonly reported visual mismatch negativity (for a review, see Stefanics et al., 2014). This finding is broadly consistent with the theory that top-down prediction signals silence matching bottom-up sensory signals and leave only the remaining prediction error to propagate forward (Friston, 2005, 2009; Rao & Ballard, 1999). Prediction also reduced the later positive P3 component (from approximately 250 ms), consistent with the theory that this component reflects involuntary orienting to novel stimuli (Friedman, Cycowicz, & Gaeta, 2001; Polich, 2007).

We also found that stimuli deviating in two feature dimensions (i.e., object deviants) evoked larger early negativities than stimuli deviating in only one feature dimension (i.e., orientation or colour deviants; *Figures 2 & 5*). This finding contradicts a previous study that found visual features elicit non-additive mismatch-related brain activity (Sulykos & Czigler, 2011), and suggests instead that the vMMN is sensitive to the extent of deviation across multiple feature dimensions. Importantly, object deviants in Sulykos & Czigler (2011) deviated in spatial frequency and orientation, whereas object deviants in the present study deviated in colour and orientation. Thus, future studies should investigate the extent to which mismatch additivity in the visual domain depends on the specific features involved.

We found that feature-based attention reduced neural responses to task-irrelevant peripheral patterns from approximately 200 ms after stimulus onset (*Figures 3 & 6*), consistent with the commonly reported ‘selection negativity’ (Gledhill et al., 2015). This effect replicated with low contrast stimuli that likely necessitated a tight focus of spatial attention (Experiment 2), contradicting the finding that feature-specific modulation of the selection negativity is contingent on spatial attention (Anllo-Vento & Hillyard, 1996; Hillyard & Münte, 1984) and suggesting instead that late effects of feature-based attention are globally effective (Gledhill et al., 2015). Interestingly, we found no difference between neural responses to congruent and incongruent stimuli earlier in the epoch (*Figures 3 & 6*), in contrast to a previous study that reported early effects of feature-based attention on neural responses to stimuli at task-irrelevant locations (beginning within 100 ms of stimulus onset; Zhang & Luck, 2009). A critical difference between Zhang & Luck (2009) and the present study is that Zhang & Luck (2009) had participants search for targets with specific feature conjunctions (luminance and colour), whereas targets in our study were defined by only a single feature (colour or orientation). Thus, it is possible that early effects of feature-based attention depend on the complexity of the attentional set. This interpretation is consistent with a recent study in which we found that neural responses to high-frequency flickering stimuli outside a search array (12.5 or 16.7 Hz, corresponding to an 80 or 60 ms cycle) are enhanced by feature-based attention during conjunction but not unique-feature search (Painter, Dux, Travis, & Mattingley, 2014).

Crucially, we found an interaction between feature-based attention and prediction in each of the two experiments. Congruent stimuli evoked smaller posterior mismatch responses than incongruent stimuli between approximately 200 and 300 ms after stimulus onset. Inspection of the ERPs revealed that the effect of feature-based attention on neural responses was larger for deviants than it was for standards. This pattern of results is consistent with our recent finding that attention enhances the processing of mismatch information from approximately 200 ms post-stimulus (Smout et al., 2019) and broadly supports the theory that attention enhances the gain of prediction errors (Feldman & Friston, 2010). Neural responses to surprising stimuli (deviants) are theorised to be modulated by attention because they contain prediction errors, whereas neural responses to predicted stimuli (standards) are less affected because they contain relatively few prediction errors. The present study extends this theory to suggest that feature-specific attentional modulation of prediction errors occurs even when the surprising stimuli are task-irrelevant and presented outside the spatial focus of attention.

Interestingly, we found that feature-based attention had no effect on the earlier vMMN evoked by deviants (109 – 160 ms). This pattern of findings contradicts a previous study that found the vMMN evoked by peripheral stimuli was smaller (more positive) when participants searched for a change in the deviating feature at fixation, relative to a different feature (Czigler & Sulykos, 2010). A subtle difference between the paradigms is that participants in Czigler and Sulykos (2010) searched for a feature ‘change’ at fixation (e.g., a change in the target object colour), whereas participants in our study searched for specific object onsets. Thus, it remains possible that subtle differences in the configuration of the attentional set can influence the timing and direction of the interaction between feature-based attention and prediction.

We manipulated target and distractor salience across the two experiments in order to investigate whether the strength of the top-down feature set modulates the neural interaction between prediction and feature-based attention. Although the pattern of neural effects did not differ between the two experiments, we observed slightly different behavioural effects as a function of task difficulty. Responses to highly salient targets (Experiment 1) that appeared immediately after a congruent pattern were slower than those to targets that appeared after an incongruent pattern. In contrast, there was no such effect of feature-congruence on responses to less salient targets (Experiment 2). These findings are broadly consistent with contingent capture theory (Folk et al., 1992), which proposes that distracting stimuli within the spatial focus of attention capture attention when they are congruent with the observers’ current attentional set. Since targets were easily detected in Experiment 1, it seems likely that some amount of spatial attention ‘leaked’ to the peripheral stimuli, facilitating contingent capture. In contrast, the higher task difficulty of Experiment 2 likely necessitated a tighter focus of attention to the central stimuli, thus prohibiting a contingent capture effect.

We did not observe an effect of predictability of peripheral patterns on target detection, or an interaction between pattern prediction and feature-congruence, in either experiment. This is consistent with a previous study that failed to find any effect of pattern prediction on response times to a central feature change target, nor an interaction with task set, at the level of single trials (though note that this study did report sustained block-wise effects on behaviour; Czigler & Sulykos, 2010). These findings suggest that the neural bias toward feature-congruent and surprising stimuli at task-irrelevant locations, observed in the present study, does not interfere with the concurrent processing of targets at task-relevant locations.

The present study contributes to a burgeoning literature on the relationship between prediction and attention. Whereas some studies have found an interaction between prediction and attention (Auksztulewicz & Friston, 2015; Jiang et al., 2013; Kok, Rahnev, et al., 2012; Marzecová et al., 2017; Smout et al., 2019), many others have reported only independent main effects (e.g. Garrido, Rowe, Halász, & Mattingley, 2017; Hsu, Hämäläinen, & Waszak, 2014; Kok, Jehee, & de Lange, 2012). We note that investigations to date have employed a wide variety of attention manipulations (e.g., feature-based, spatial, temporal) and prediction manipulations (e.g., first-order, rule-based) across different sensory modalities (e.g., visual, auditory). Thus, the equivocal pattern of findings to date may stem from distinct relationships between different subprocesses of attention and prediction across the various modalities. In particular, previous studies that found an interaction between visual attention and prediction presented stimuli at attended locations (Jiang et al., 2013; Kok, Rahnev, et al., 2012; Marzecová et al., 2017; Smout et al., 2019) or used paradigms that did not require focussed attention to complete the task (Czigler & Sulykos, 2010), leaving open the possibility that spatial attention is necessary in the interaction with prediction. The present study extends this literature by demonstrating that visual predictions interact with feature-based attention to modulate neural responses to stimuli outside the spatial focus of attention. The nature of this interaction is consistent with the theory that attention optimises the expected precision of predictions by modulating the gain of prediction errors (Feldman & Friston, 2010). Future research should continue to parse ‘attention’ and ‘prediction’ into more precise taxonomies that reflect specific mechanisms in the brain and investigate potential interactions between each of these subcomponents. This work could illuminate the extent to which predictive coding theory might be considered a ‘unified theory of the brain’ (Friston, 2010).

